# Assessing the functional impact of protein binding site definition

**DOI:** 10.1101/2023.01.26.525812

**Authors:** Prithviraj Nandigrami, Andras Fiser

## Abstract

Many biomedical applications, such as classification of binding specificities or bioengineering, depend on the accurate definition of protein binding interfaces. Depending on the choice of method used, substantially different sets of residues can be classified as belonging to the interface of a protein. A typical approach used to verify these definitions is to mutate residues and measure the impact of these changes on binding. Besides the lack of exhaustive data this approach generates, it also suffers from the fundamental problem that a mutation introduces an unknown amount of alteration into an interface, which potentially alters the binding characteristics of the interface. In this study we explore the impact of alternative binding site definitions on the ability of a protein to recognize its cognate ligand using a pharmacophore approach, which does not affect the interface. The study also provides guidance on the minimum expected accuracy of interface definition that is required to capture the biological function of a protein.

**AUTHOR SUMMARY:** The residue level description or prediction of protein interfaces is a critical input for protein engineering and classification of function. However, different parametrizations of the same methods and especially alternative methods used to define the interface of a protein can return substantially different sets of residues. Typical experimental or computational methods employ mutational studies to verify interface definitions, but all these approaches inherently suffer from the problem that in order to probe the importance of any one position of an interface, an unknown amount of alteration is introduced into the very interface being studied. In this work, we employ a pharmacophore-based approach to computationally explore the consequences of defining alternative binding sites. The pharmacophore generates a hypothesis for the complementary protein binding interface, which then can be used in a search to identify the corresponding ligand from a library of candidates. The accurate ranking of cognate ligands can inform us about the biological accuracy of the interface definition. This study also provides a guideline about the minimum required accuracy of protein interface definitions that still provides a statistically significant recognition of cognate ligands above random expectation, which in turn sets a minimum expectation for interface prediction methods.

## INTRODUCTION

Protein-protein interactions are key players in many biological processes including metabolism, development, and regulation. Residue level descriptions of protein binding interfaces are essential for explaining, classifying, and modulating the formation of specific protein complexes. The biological function of a protein binding interface critically relies on its specificity, the ability to selectively recognize its cognate binding partners. However, the question of what exactly comprises a biological interface does not have a clear answer. Many interface prediction approaches exist. Some of these employ a radial distance cutoff-based approach (1, 2), which can be combined with a requirement of compatibility of interactions (3), while others use Voronoi-polyhedra based calculations, such as INTERCAAT (4), QCONS (5) and others (6, 7). Another group of methods focus on monitoring changes in the accessible surface area upon complex formation, such as NACCESS (8, 9) or POPSCOMB (10), which are all based on the original method of Lee and Richards (11). However, there is a lack of consensus among these methodologies regarding how to consistently define the biologically relevant protein-protein interaction interface (12–19). A recent study illustrated that even among approaches employing nearly identical algorithms to define interfaces in known protein complexes, a minimal difference in definition can reduce the agreement between them to about 80% and in a significant number of cases the interface definitions could overlap by as little as 40% (20). The differences of interface definitions among different approaches are even larger. From another point of view, it is important to know what level of interface definition inaccuracy is acceptable while still maintaining a useful prediction. Given the uncertainty in defining a protein interface, it would be important to know how much one can mispredict or misdefine the cognate biological interface and still capture its biological function, i.e., to successfully predict or identify an interface that is sufficient to selectively recognize its cognate partner.

A major strategy, both computationally (21–25) and experimentally (26–30) used to verify a biologically relevant interface is alanine scanning mutagenesis. Another similar but complementary strategy is to mutate interface positions with closely related amino acids (31, 32). However, the availability of complete datasets that exhaustively explore the interface of a given protein by measuring the impact of mutations on binding affinity is limited (5). Besides this practical limitation, a more important, conceptual problem is that the effect of mutation on the overall stability of a receptor-ligand complex is not straightforward to address, in part, due to the conformational alterations associated with induced mutations (33). An additional consideration is that positions outside of the binding interface, that do not play a role in the direct recognition of the cognate ligand, can have an indirect effect on recognition if altered. Mutations at distal sites from the interface can either increase (34, 35) or diminish the binding affinity (36) of a receptor against a ligand. All these computational and experimental approaches inherently suffer from the problem that in order to probe the importance of any one position of an interface, an unknown amount of alteration is introduced into the very interface being studied. Taking this into account, one can conclude that it is not straightforward to define the binding interface of proteins by probing the impact of residue mutations, which are often associated with global structural modulation of the protein (37).

In this paper, we computationally explore various protein interface definitions by probing the minimum interface necessary to successfully recognize its cognate ligand. Through this analysis we quantitatively investigate how much overlap alternative receptor interface patches must share with their true biological interface to still recognize their cognate partner reliably. Similarly, we investigate how many true interface residues can be lost without diminishing a proteins ability to accurately recognize its cognate partners.

The analysis presented here does not perturb the integrity of the protein structure with mutations. Instead, we utilize a computational approach, ProtLID (38), to obtain a residue-specific pharmacophore (rs-pharmacophore) description for the protein interface. A pharmacophore is an abstract description of the critical atoms, groups, charged regions, and their spatial distributions that are essential for the biological activity of a small-molecule drug (39, 40). The preferred, complementary residue positions on the receptor interface are predicted via rigorous molecular dynamic sampling of single residue probes. The consolidated spatial preferences establish a unique fingerprint of the single-residue probe preferences on a hypothetical ligand interface. The rs-pharmacophore is then used to screen candidate ligands for the given protein receptor where each candidate ligand is ranked according to the degree of match against the predicted rs-pharmacophore. The candidates that are highly ranked are the predicted binding partner(s) of the given protein (41). ProtLID was successfully used to identify the cognate partners for a given receptor (38), and to redesign various protein interfaces for ligand binding specificity (42, 43). By employing ProtLID, we do not introduce any alterations into the protein but instead construct complementary rs-pharmacophore descriptions for each alternatively defined binding interface. These alternative interfaces are then tested against a library of candidate ligands to see which alternatively defined interfaces retain their ability to recognize their cognate ligands from a set of decoys.

In this work, we study three protein interfaces: PD1:PD-L1, CTLA4:CD80, and CTLA4:CD86. These interactions are critical in regulating cellular functions. PD1 and PD-L1 form a co-inhibitory complex that can limit the development of T-cell response. PD1-PD-L1 interaction helps ensure that the immune system is activated only at the appropriate time to minimize autoimmune inflammation (44). CTLA4, on the other hand, is the first immune checkpoint receptor to be clinically targeted. It is expressed exclusively on T cells where it primarily regulates the amplitude of the early stages of T cell activation (45).

Our results show that, on average, one can misdefine a protein interface by about 20-30% while still preserving its ability to recognize its cognate partner with statistically significant results. These results suggest that methods that predict protein interfaces should achieve above 70% accuracy to be useful. Our results also show that receptors with higher binding affinity (CTLA4:CD80) compared to receptors with lower binding affinity (CTLA4:CD86) are harder to destroy by removing true interface residues and adding false interface residues.

## RESULTS

### Generating and assessing alternative interface definitions of the same size

The goal of this work is to assess the impact of alternative interface definitions for a protein receptor by monitoring how well each alternatively defined interface can recognize its cognate binding partner(s) from an ensemble of candidates. We consider the receptor-ligand interactions in three protein-protein complexes: PD1:PD-L1, where the interface on PD1 comprises 14 residues, and CTLA4:CD86/CD80, with a smaller interface on CTLA4 of 9 and 10 residues, respectively (Fig 2). CTLA4 was selected because ProtLID was shown to accurately assign a high ranking to its two known cognate ligands, while PD1 was selected because it has many known cognate ligands, which can provide statistical power for the observations, although with somewhat lower accuracy. First, we generated a reasonably large number of alternative poses of the cognate protein-protein complex in question. From those poses, alternative interface patches were selected with the same interface size as, but with varying degrees of overlap with, the original interface. The number of residues in the interfaces was kept constant in order to avoid dealing with the confounding impact of varying interface sizes when recognizing the cognate ligand. To sample physically plausible alternative interface patches, we employed the docking software ZDOCK (46) to generate 2000 top scoring docked poses by keeping the receptor PD1 or CTLA4 fixed, and allowing their biological binding partners, PD-L1, CD80 or CD86 to dock on the surface of the receptor. The interfaces of the receptor in all the cognate poses were determined using the program CSU (3). From the 2000 conformations that ZDOCK generated, we extracted all complexes with interface patch sizes equal to the original interface and varying degree of overlap with the original interface on the receptor protein (Fig. 1). In case of PD1:PD-L1, out of the 2000 docked complexes 181 poses had the same number of residues (14) as the original interface, for CTLA4:CD80 and CTLA4:CD86, 230 and 255 such complexes were found, respectively. For PD1:PD-L1, CTLA4:CD80, and CTLA4:CD86, out of the 181, 230, and 255 cases 99, 44, and 64 cases had no overlap with the original interface at all, respectively (Fig. 3).

**Figure 1:**
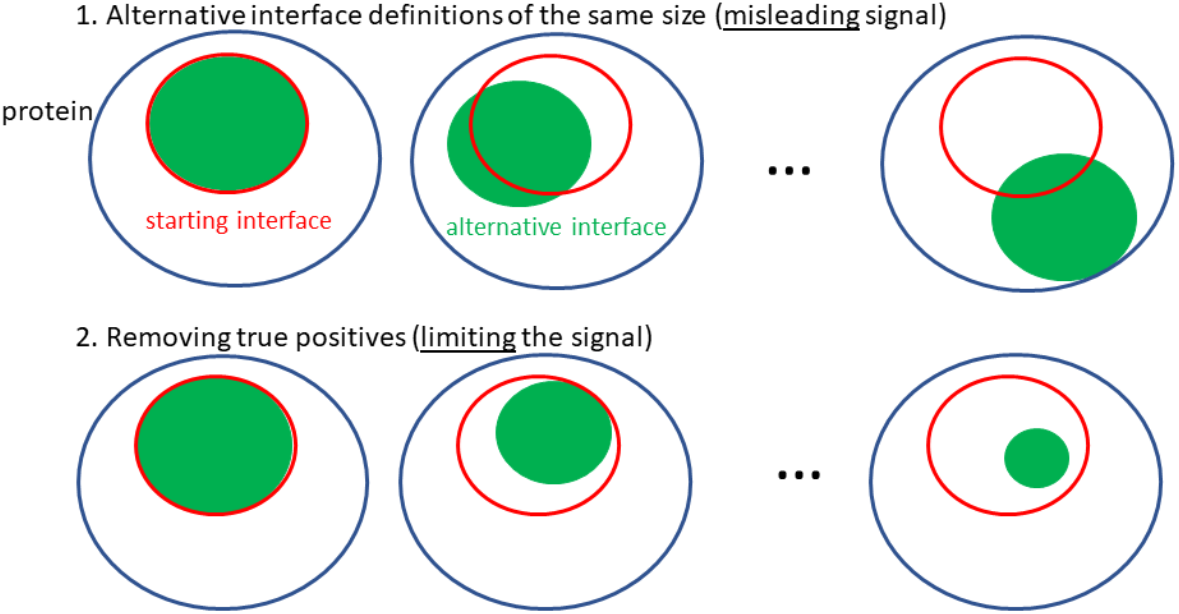
Schematic representation of different alternative interfaces explored in this work. In the first approach, each alternative interface generated preserves the same number of residues as the starting interface but with varying degrees of overlap with it. In the second approach, residues from the original interface are removed combinatorically.

**Figure 2:**
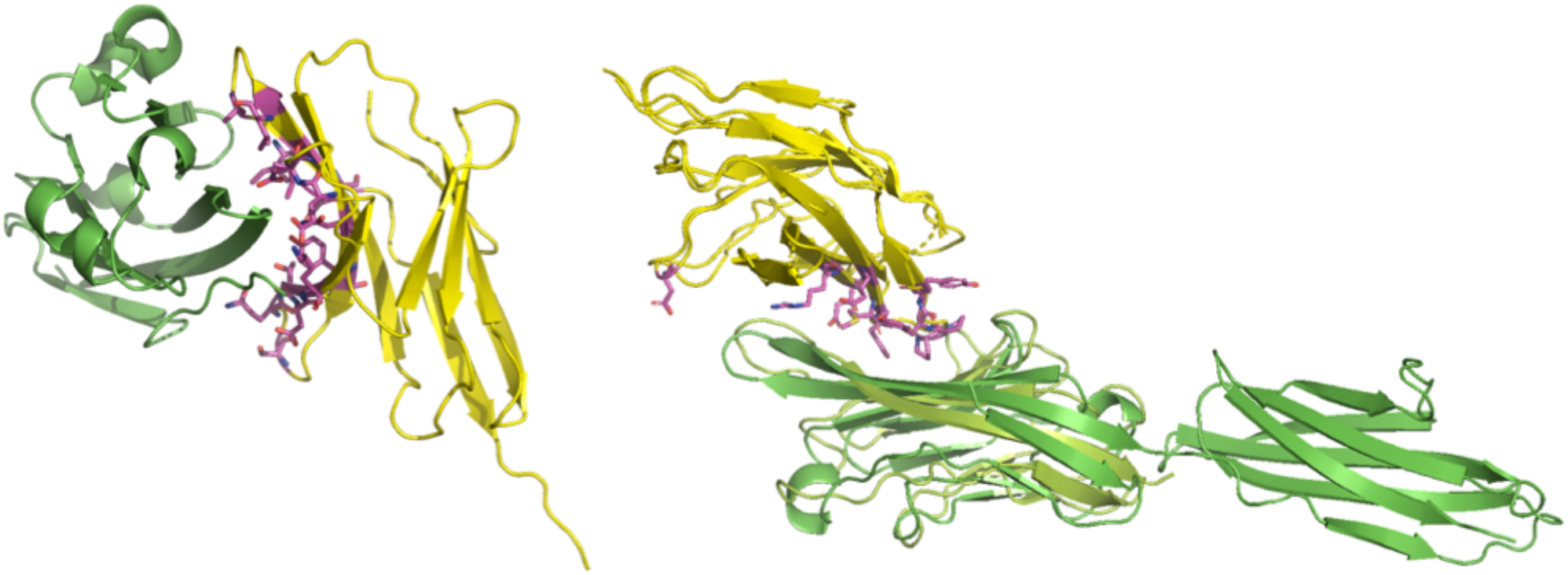
PD1-PDL1 and CTLA4-CD80 cognate complexes. PD1 is shown in yellow and PDL1 is shown in green (left). CTLA4 is shown in yellow, CD80 is shown in dark green, and CD86 is shown in light green (right). Interface residues on CTLA4 (for CTLA4-CD80 complex) are shown as sticks. Visualization rendered in Pymol.

**Figure 3:**
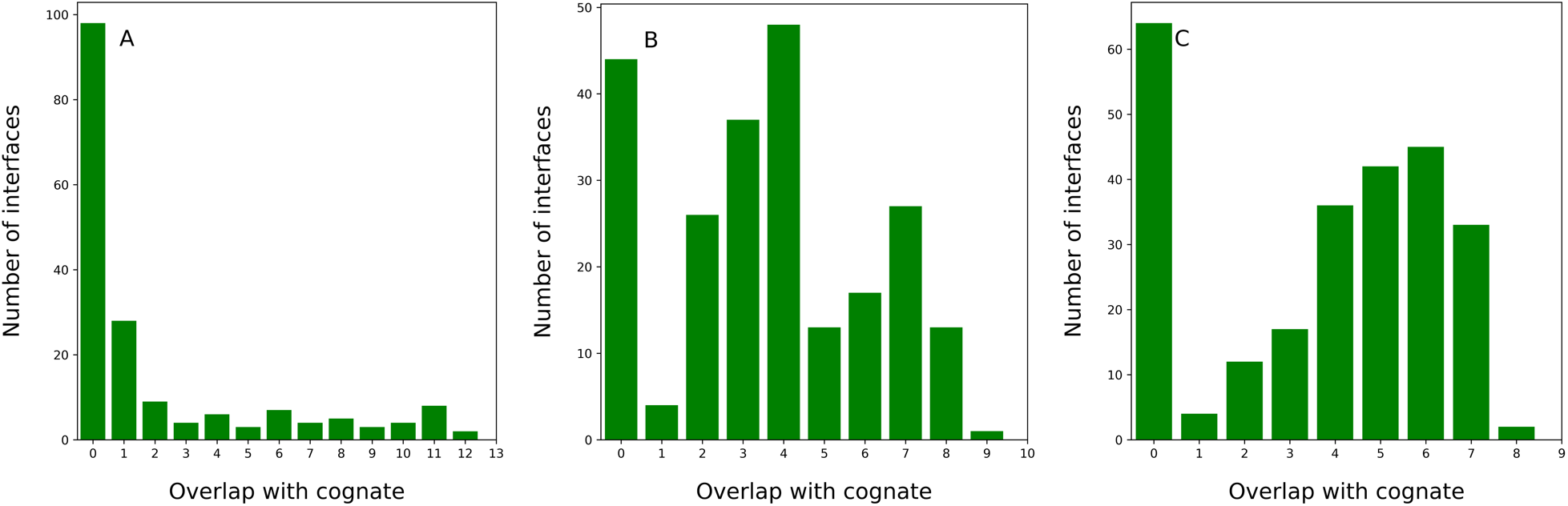
Distribution of docked poses comprising the same number of residues as the cognate protein-protein complex for (A) PD1-PDL1, (B) CTLA4-CD80, and (C) CTLA4-CD86 interfaces, generated by ZDOCK. The X- axis shows the number of residues from each alternative pose overlapping with the original interface.

For each alternative interface definition, a new rs-pharmacophore was calculated, and a ligand search was performed to assess the accuracy of the pharmacophore. In other words, we tested how well each alternative pharmacophore was able to identify its cognate ligand from a library of 103 IgSF structures (Suppl. Table 1). Fig. 4 shows the number of cognate ligands identified in the top 20^th^ percentile for the PD1 receptor when we consider interface patches with gradually reduced degree of overlap with the original interface. The original interface (14 out of 14 residues overlap with the original interface (Fig. 4)), was able to capture 13 out of 24 ligands in the top 20% of all candidate ligands screened. None of the alternatively defined interfaces provided by ZDOCK surpassed this performance, which suggests that the original interface definition was the best available one. As we considered alternative interfaces with reduced numbers of residual overlaps with the original interface, a gradual drop in accuracy was observed, with fewer and fewer cognate ligands recognized in the top 20%. It appears that approximately 70% of the original interface needs to be correctly captured to reach a signal that is statistically significantly distinguishable from random ranking (Fig. 4). If we consider only the rs-pharmacophore results that were reliable (based on the skewness of the distribution of ProtLID scores), they performed better on average than considering signal from both reliable and unreliable interface patches. This is shown by the higher levels of black data points compared to the blue dotted line. Statistically, if the 24 cognate ligands with known structures were uniformly distributed (i.e., randomly predicted) among the ligand candidates, then one would expect 20% of these to be identified by chance in the top 20% of all ligands. This theoretically expected level is marked by a horizontal orange line at approximately 4.67 in Fig. 4 for reference. To test empirically this hypothetical expectation, we randomly selected 10 interface patches from ZDOCK poses, which had no overlap with the original interface at all (0 on x-axis) and tested how the corresponding pharmacophores recognized their cognate ligands. First, all predictions were correctly identified as not reliable (empty black circles), second, the average of these rankings showed a faithful reproduction of recognizing about 4.67 ligands as expected by random chance. The standard error of the mean of the resulting signal for each cohort of patches shows a narrower uncertainty when the number of constituent patches analyzed is higher, and the range remains reasonably above the random expectation until about 70% of the original interface is captured accurately.

**Figure 4:**
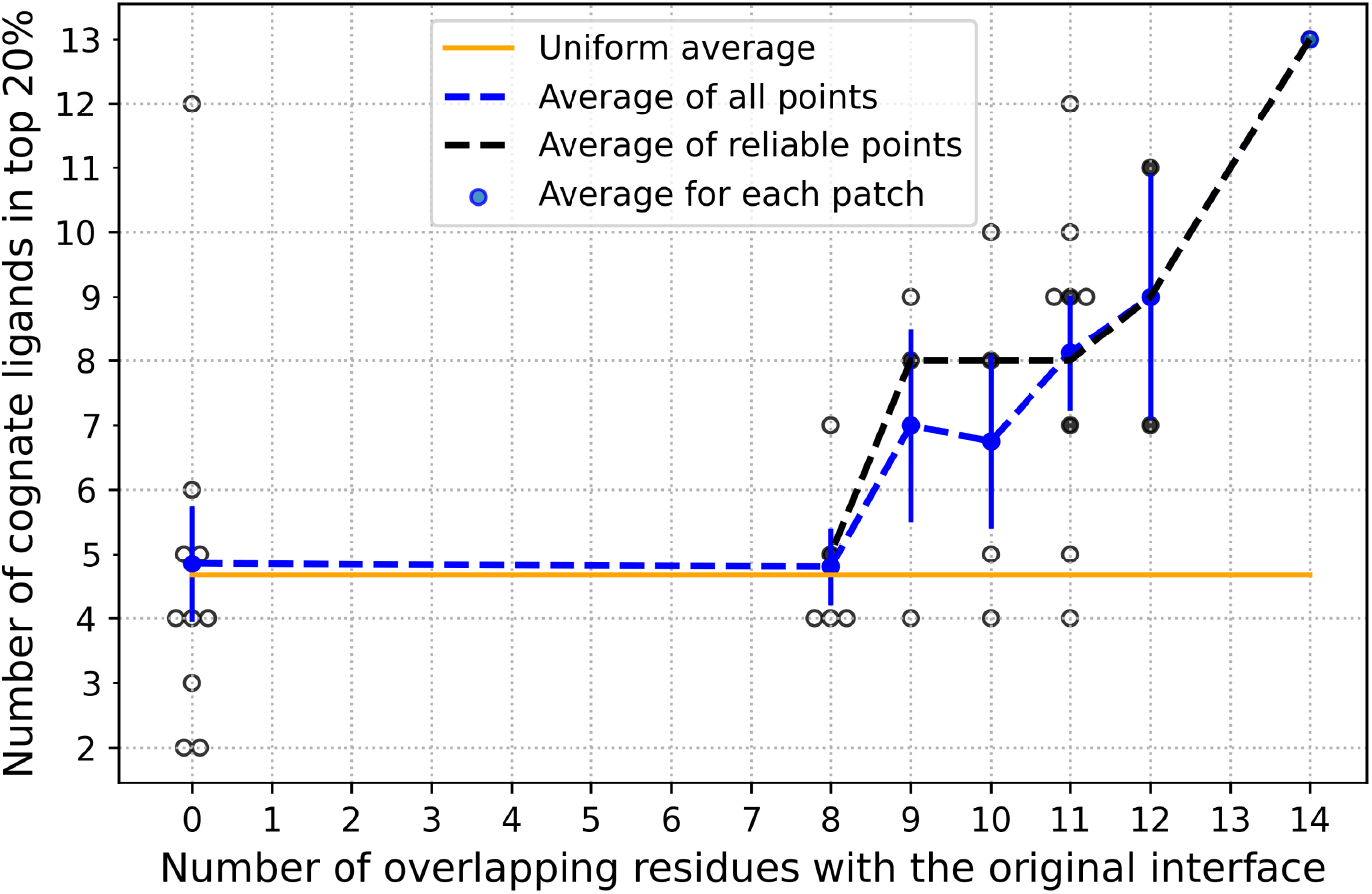
Number of cognate ligands ranked in the top 20 percentile as we deviate away from the original interface of PD1:PDL1. The number of residues overlapping between the alternative and the original interface is shown on the X-axis. Solid black circles are reliable and hollow circles are unreliable predictions. The blue circles and corresponding blue dotted line shows the average number of ligands recognized for all receptor patches explored, while black dotted line connects average results for reliable prediction only. The orange line marks a theoretically expected random reference assuming uniform ranking of ligands, which is 4.67. The vertical blue lines show the standard error of the mean.

It is interesting to consider protein receptor interfaces with a smaller number of known cognate ligands as well. We consider two CTLA4 interfaces in complex with CD80 and CD86. We collected all the docked poses of CTLA4:CD80 and CTLA4:CD86 generated by ZDOCK that have the same interface size as CTLA4 in complex with its respective cognate binding partners (Fig. 2). Because CTLA4 only has 2 cognate ligands, we did not monitor the fraction of the ligands found in the top 20% of all hits as we did before. Instead, we directly monitored the actual average ranking of the CTLA4 cognate ligands (Fig. 5). Once again, cognate ligand rankings were computationally calculated on several interface patches by systematically reducing the degree of overlap with the true cognate interface residues, while keeping the total number of interface residues constant. Like we observed with the PD1:PD-L1 complexes, the interface reliability tends to disappear if more than 20-30% of residues from the original interface is lost. Finally, as before, none of the alternatively defined interfaces produced higher accuracy results than the original interface. This may simply be a consequence of the fact that the cases we picked generally performed well in the original testing to rank cognate ligands, which requires that the interfaces had to be correctly defined. We picked better performing cases to start with because we wanted to monitor how the ability of an interface to distinguish its cognate binding partners decreased as the accuracy of its interface definition decreased.

**Figure 5:**
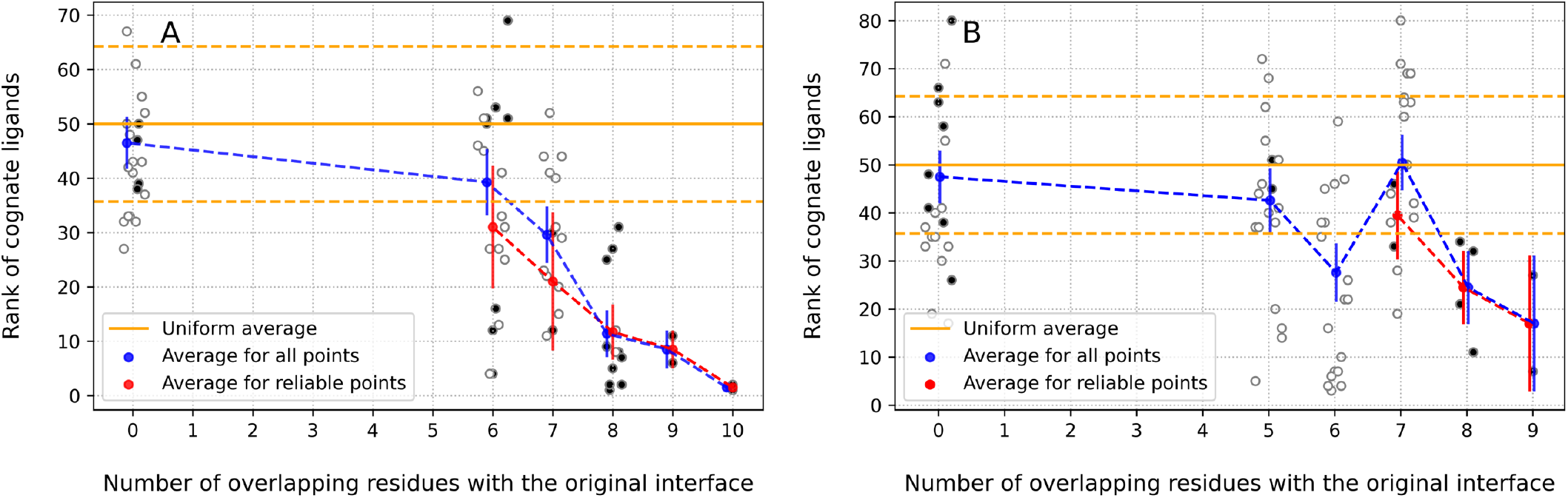
The percentile rank of cognate ligands (Y-axis) as a function of overlap between the alternative interface and the original interface. Solid black circles are reliable and hollow circles are unreliable predictions. (A) The average percentile rank of cognate ligands as we deviate away from the original interface of CTLA4:CD80, and (B) in CTLA4:CD86 interface. Each ProtLID result is assessed for reliability: solid circles are reliable and hollow circles are unreliable. Blue circles show the average results for a given interface size for all interfaces, while pink circles show the average of only the reliable predictions. The corresponding standard error of the mean around the averages of all points and only reliable points are shown by vertical blue and pink lines, respectively. The solid orange line shows the random expectation, with the corresponding standard error of the mean shown by the orange dashed lines.

The large degree of overlap between the two CTLA4 interfaces (eight residues out of 9 and 10, respectively), defined using the CTLA4:CD80 complex or the CTLA4:CD86 complex, enables us to assess the ability of the CTLA4:CD80 interface to recognize CD86, and the CTLA4:CD86 interface to recognize CD80. As shown in Fig. S1, the ability of the CTLA4:CD80 interface to recognize CD86 as a cognate ligand is poorer than the average ability of it to recognize its own cognate ligands. In contrast, as shown in Fig. S2, the ability of the CTLA4:CD86 interface to recognize CD80 as a cognate ligand is better than the average ability of it to recognize its own cognate ligands. This suggests that CD80 has greater specificity than CD86 for recognizing the CTLA4 interface.

In general, we find that the performance of the CTLA4:CD80 interface is better on average than that of the CTLA4:CD86 interface in recognizing cognate partners. This observation possibly implies that the binding affinity of CTLA4 for CD80 is markedly higher than the affinity for CD86 (47–49). The CTLA4:CD80 interface recognizes its two known cognate partners as ranks 1 and 2 while the CTLA4:CD86 interface recognizes its two known cognate binding partners as ranks 7 and 27.

### Removing true positive residues from the interface

In an alternative approach, we explored what happens when we gradually eliminate residues in a combinatorial way from the original interface, without replacing them with non-interface residues (thereby reducing the size of the interface). We systematically removed residues in cohorts (for example, singles, pairs, triplets etc.) from the residues comprising the rs-pharmacophore to assess the ability of the reduced rs-pharmacophore to recognize its cognate ligands. Fig. 6 shows the ability of PD1 to recognize its cognate ligands. Two assessment metrics are used – the average number of cognate ligands in the top 20 percentile identified by a given pharmacophore and the overall average rank of cognate ligands identified by a given pharmacophore. First, considering only those data points that are predicted to be reliable produces an improvement of the output signal. Second, when approximately 78% of the original interface residues remain in the pharmacophore, the average rank of cognate ligands and the average number of cognate ligands in the top 20 percentile approaches the random limit. Once again, we observe that the best performance in recognizing cognate ligands is observed for the pharmacophore generated from the original interface. Next, we repeated the same exercise with CTLA4 in complex with CD80 and CTLA4 in complex with CD86. Since there are only two cognate ligands for CTLA4 with known crystal structures, we used the metric of the average percentile rank of cognate ligands. One key difference between the performance of two systems (Fig. 7) is that the CTLA4:CD80 interface results in a substantially stronger ranking for the cognate ligands compared to the CTLA4:CD86 interface. In addition, it appears that it is somewhat more difficult to “destroy” the cognate ligand recognition ability of the CTLA4:CD80 interface compared to the CTLA4:CD86 interface. Even at a 60% overlap, the CTLA4:CD80 interface recognizes its cognate ligands at a statistically significant level. Like the PD1:PD-L1 interface, the results show that, on average, the output signal improves by considering only the reliable data points. When approximately 60-78% of the true interface residues remain in the pharmacophore, the average rank of cognate ligands approaches the random limit. None of the alternative interfaces outperformed the original one, i.e., initial receptor interfaces studied in this work faithfully represent a reasonably accurate description of the true biological interface because these show the best performance in recognizing their corresponding cognate ligands.

**Figure 6:**
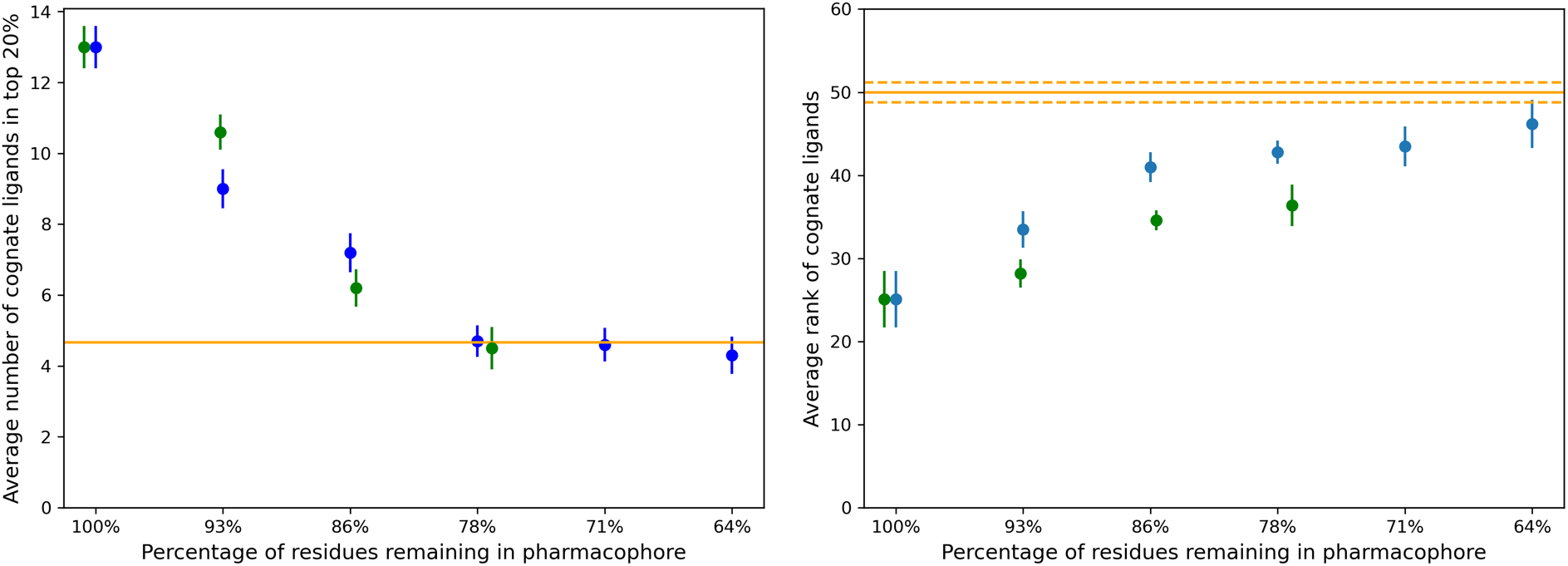
Assessment of PD1 ability to recognize its cognate ligands as we gradually reduce the number of residues from the original receptor interface. (Left) Average number of cognate ligands in top 20 percentile for all (blue) and reliable (green) data points. (Right) Average rank of cognate ligands for all (blue) and reliable (green) data points with the corresponding standard error of the mean shown by vertical lines. Orange solid line represent the random expectation, with the orange dashed line showing the corresponding standard error of the mean: (left) the average number of cognate ligands in the top 20% is about 4.67 and (right) average rank of all ligands is about 50%.

**Figure 7:**
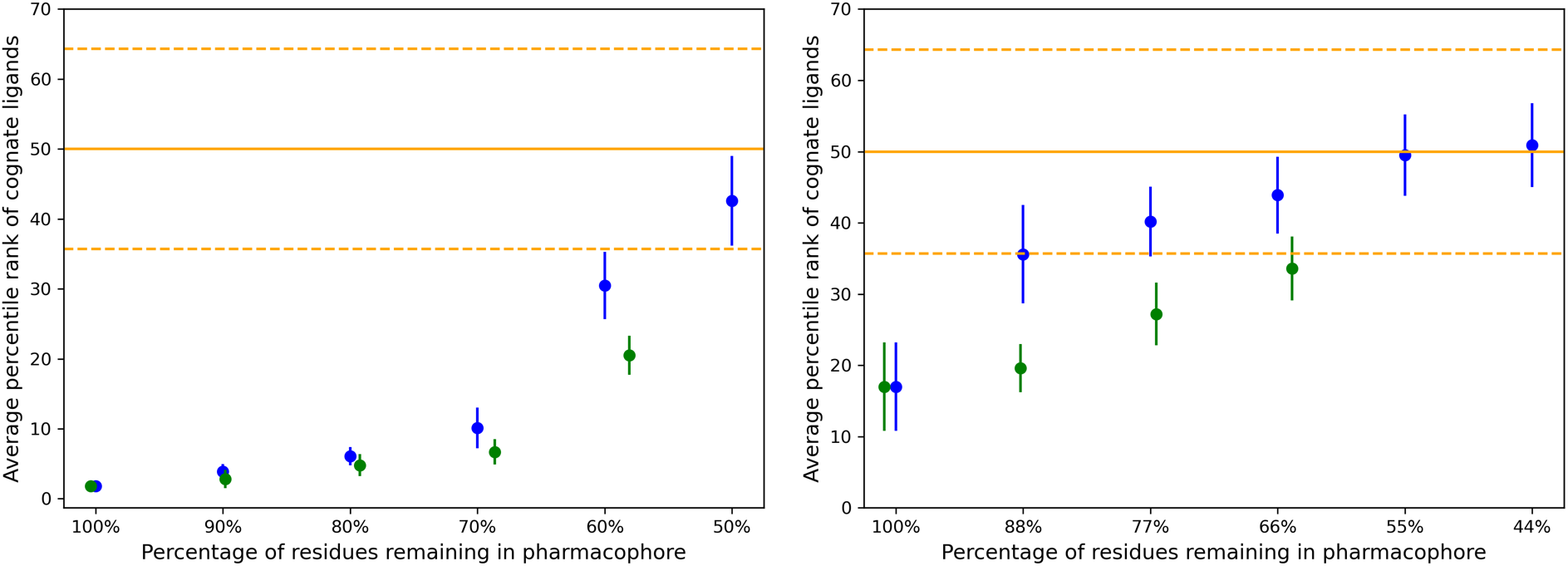
Assessment of recognizing the cognate ligand (CD80 (left) and CD86 (right)) of CTLA4 receptor interface. Average percentile rank of cognate ligands is shown (Y-axis) as a function of gradually reduced interfaces (X-axis). Blue and green points represent all and only reliable data points, respectively with the corresponding standard error of the mean shown by vertical lines. The orange solid line shows the random expectations for average rank of cognate ligands, and orange dashed line shows the standard deviation.

## DISCUSSION

Protein interfaces can be defined using various approaches, which include using radial cutoffs, monitoring change in solvent accessible surface areas, or employing Voronoi polyhedra calculations. The disagreement among the results of these methods can be traced back to the fact that these approaches aim at identifying interfaces from a physical point of view using alternative criteria. In contrast, in this work we tried to assess the impact of alternative interface definitions from a biological point of view, from the perspective of a protein to maintain its ability to selectively recognize its cognate binding partner. The central question we ask in this work is how imperfect can the interface definition be whilst retaining the ability to recognize the cognate ligand(s). We assessed this by pursuing alternative interface patches that had decreasing overlap with the original biological interface. In this work, none of the alternative interfaces sampled produced higher accuracy results than the original definition obtained from CSU. This does not validate CSU as a uniformly accurate approach, but simply reflects the fact that we picked well performing cases, where the pharmacophore reliably identified the cognate ligand partners from a set of decoys. This means that the original definition of the interface had to be reasonably good to start with.

The results show that on average, we can safely lose approximately 20-30% of the true biological interface and still recognize the cognate ligands of the receptor with reasonable, albeit lower, confidence than the original interface. Additionally, we observe that the skewness of ProtLID scores is informative to identify reliable alternative interface definitions. These also results provide guidance for interface prediction methods. Current methods fall in two major categories, *ab-initio* methods (50, 51) and template-based methods (20). *Ab-initio* methods are more broadly applicable but their accuracy ranges between 30-40%, while template-based method can reach higher prediction accuracies, but they rely on the availability of known template (20, 52). A recent combined approach reported F-score accuracies just above 0.5 (52). Meanwhile the results from the current study suggest that prediction methods should really reach F-score ~0.7 level to produce interface predictions that are biologically useful for practical applications and highlights the need to develop methods that breach this performance gap.

## METHODS

### Protein interfaces

We study three protein-protein heterodimer complexes in this work, PD1:PD-L1 (PDB code: 4ZQK), CTLA4:CD80 (PDB code: 1I8L), and CTLA4:CD86 (PDB code: 1I85). A rs-pharmacophore is calculated with ProtLID treating PD1 (4ZQK, chain B), CTLA4 (1I8L, chain C), and CTLA4 (1I85, chain D) as receptor proteins. Each receptor rs-pharmacophore is screened against a library of 103 structures of the immunoglobulin superfamily (Suppl. Table 1). There are 24 available structures of the cognate ligand of receptor PD1 (Protein Data Bank (53) codes: 4ZQK.A, 3BIK.A, 3BIS.A, 3FN3.A, 3SBW.A, 4Z18.A, 5C3T.A, 5GGT.A, 5GRJ.A, 5IUS.C, 5J89.A, 5J8O.A, 5JDR.A, 5JDS.A, 5N2D.A, 5N2F.A, 5NIU.A, 5NIX.A, 5O45.A, 5O4Y.B, 5X8L.A, 5X8M.A, 5XJ4.A, 5XXY.A). On the other hand, CTLA4 has 2 cognate binding partners with known structure: CD80 (1I8L.A and 1DR9.A) and CD86 (1I85.B and 1NCN.A).

### Docking

ZDOCK (46) was used to perform rigid body docking. In each case the receptor was kept rigid, and the ligand was docked onto the receptor surface to generate 2000 top scoring alternative poses of the receptor-ligand complex.

### Interface definition

CSU (Contacts of Structural Units) (3) was used to determine the interacting residues based on interatomic contacts distances (using the standard 4.5 Å cutoff) and complementarity of interacting atomic groups in the complex structures.

### ProtLID

Protein Ligand Interface Design, (38) is a computational method that generates a rs-pharmacophore description for a given protein interface. By running extensive molecular dynamics simulations of single-residue probes, ProtLID calculates the optimal complementary interface. The resulting residue-based pharmacophore (rs-pharmacophore) comprises the residue types and location preferences on the complementary interface. This rs-pharmacophore is subsequently used to find potential matches among candidate ligands using a pattern matching algorithm (41).

### Assessing the reliability of ProtLID pharmacophore

Mathematical skewness of ProtLID scores assesses the reliability of the rs-pharmacophore generated by ProtLID (38). Skewness is defined as, *skewness* = (*Pm3/ Psd^3^*) where *Pm3* is the third moment of a ProtLID score distribution, and *Psd* is the standard deviation of ProtLID scores. Once a pharmacophore for a given protein interface is generated, we enumerate all possible 5-mer combinations of the calculated residue preferences to screen a ligand structure database. This results in a certain number of matches for each potential ligand out of all combinatorial possibilities. For instance, for a 15-residue large rs-pharmacophore, the number of 5-mer enumerations is 3003 (15C5). The ProtLID score is the number of 5-mer hits for a particular ligand. Skewness is calculated over the distribution or scores obtained for all possible ligands. Only interface patches for which the skewness is above 2.5 are deemed reliable (54), while others are unreliable.

## ACKNOWLEDGMENTS

This work was supported by National Institutes of Health (NIH) grants GM136357 and AI141816.

## SUPPORTING INFORMATION CAPTIONS

Figure S1: The percentile rank of cognate ligands as a function of overlap of the alternative interface definitions with the original interface of CTLA4:CD80. Solid black circles are reliable and hollow circles are unreliable predictions. Starting with the CTLA4:CD80 interface, we calculate the average and standard errors of ranking CD86 ligand (green circles and green vertical lines). The number of residues common between CTLA4:CD80 and CTLA4:CD86 interfaces is 8. Green and blue lines correspond to reliable and all data points, respectively. The orange solid line and dashed lines show the random expectation and corresponding standard error of the mean, respectively.

Figure S2. As in Fig S1, but we explored the ranking of CD80 for the alternative CTLA4:CD86 interface definitions.

Table S1: List of 103 decoy structures used to screen rs-pharmacophore for the protein receptors studied.

